# Maternal control over caste allocation in the ant *Cardiocondyla obscurior*

**DOI:** 10.1101/2021.08.13.456267

**Authors:** Eva Schultner, Tobias Wallner, Benjamin Dofka, Jeanne Bruelhart, Jürgen Heinze, Dalial Freitak, Tamara Pokorny, Jan Oettler

## Abstract

Ant queens and workers commonly engage in conflict over reproductive allocation because they have different fitness optima. How such conflicts are resolved depends on the power that each party of interest holds. Here, we show that queens have control over caste allocation in the ant *Cardiocondyla obscurior*. With the discovery of crystalline deposits that allow the identification of queens and workers across the entire course of development, we demonstrate that caste fate is irreversibly determined in the egg stage. Comparison of queen and worker-destined eggs and larvae revealed size and weight differences in late development, but no discernible differences in traits that may be used in social interactions, including hair morphology and cuticular odors. In line with this lack of caste-specific traits, adult workers treated developing queens and workers indiscriminately, even in the presence of a fertile queen. Together with previous studies showing queen control over sex allocation, these results provide evidence that conflict over reproductive allocation is absent in this ant.

## Introduction

Ant queens and workers are fixed in irreversible, evolutionarily co-dependent roles, analogous to the germ line and soma of multicellular organisms [1–3]. However, compared to multicellular organisms, ant colonies can show relatively high levels of within-unit conflict. This is because they are made up of non-clonal individuals, with factors such as polygyny (multiple reproducing queens per colony) and polyandry (multiple mating by queens) causing variation in relatedness among nestmates [4,5]. As a result, colony members can differ in their fitness optima regarding reproductive allocation, creating the potential for conflict [6–8].

Whether potential conflict leads to actual conflict depends on the power each party holds, in particular over reproduction and access to information [9–11]. Adult reproductive power can be classified along a range of worker reproductive constraints [12–15]: in species where workers retain fully or partly functional reproductive organs (most ant genera), adult females can engage in conflict over reproductive dominance or male parentage by participating in production of female and male offspring [16–18]. In the few ant genera in which workers have lost reproductive organs entirely, they cannot participate in reproduction directly. Nonetheless, because workers are responsible for brood care, even in these species they can influence reproductive allocation by manipulating the sex and caste ratios of brood, e.g by preferentially rearing females into queens or by culling male or queen-destined brood [19–22]. Across the range of reproductive constraints, the fitness interests of developing individuals also play an important role, in particular in the context of queen-worker caste determination [21,23,24]. In all conflict scenarios, access to information about brood maternity, sex and caste can be decisive, placing developing individuals at the center of within-colony conflict [25,26].

*Cardiocondyla* ants represent the extreme end of the range of reproductive constraints because workers lack reproductive organs, precluding any direct conflict over reproduction. In *Cardiocondyla obscurior*, workers further refrain from manipulating brood sex ratios, giving queens control over colony-level sex allocation [27,28]. Recently, we found that in single queen colonies of this ant, production of new queens increases as the resident queen ages, irrespective of absolute queen fitness, queen lifespan or colony size [29]. This suggests that queens also control caste allocation, and points towards relaxed selection on potentially costly traits related to caste allocation conflict.

Here, we explore this idea by providing a description of the phenotypes of developing *C. obscurior* ant queens and workers, made possible by the discovery of crystalline deposits that distinguish castes from the egg stage onwards. We focus on traits that are predicted to differ between castes and could transmit potential information about caste in a social context, and test whether workers discriminate between developing queens and workers. Our results support the hypothesis that queens control primary caste ratios in eggs and workers refrain from manipulating secondary caste ratios of brood, giving queens complete control over caste allocation in this ant.

## Results and Discussion

### Identifying female caste in developing ants

“Auch ein blindes Huhn findet mal ein Korn” Twenty years after the first *C. obscurior* colony was brought to the laboratory in Regensburg, we discovered that queen and worker-destined eggs and larvae can be distinguished by the caste-specific distribution of white spots in the posterior end (Figures 1,2). To our knowledge, this is the first description of a discrete trait that can be used to identify female caste over the entire course of ant development. Semi-thin sections of third instar larvae viewed through a polarization filter revealed these spots have a crystalline reflection. Queens exhibit a pair of clumped crystalline deposits on the dorsal side, which become larger and more distinct as development progresses (Figures 1,2, Figure S1). In late-instar queen larvae, crystalline deposits form large aggregations around the developing ovaries, which already contain immature oocytes (Figure 3). Workers do not have crystalline deposits in the egg stage (Figure 2), and small, randomly distributed deposits in larval stages (Figures 1, 3). In both castes, crystalline deposits may consist of urate crystals, which are usually stored in specialized cells, the urocytes, in the insect fat body [30], and which have been linked to nutrient storage in ant larvae [31] and oocyte development in termite reproductives [32].

**Figure 1:**
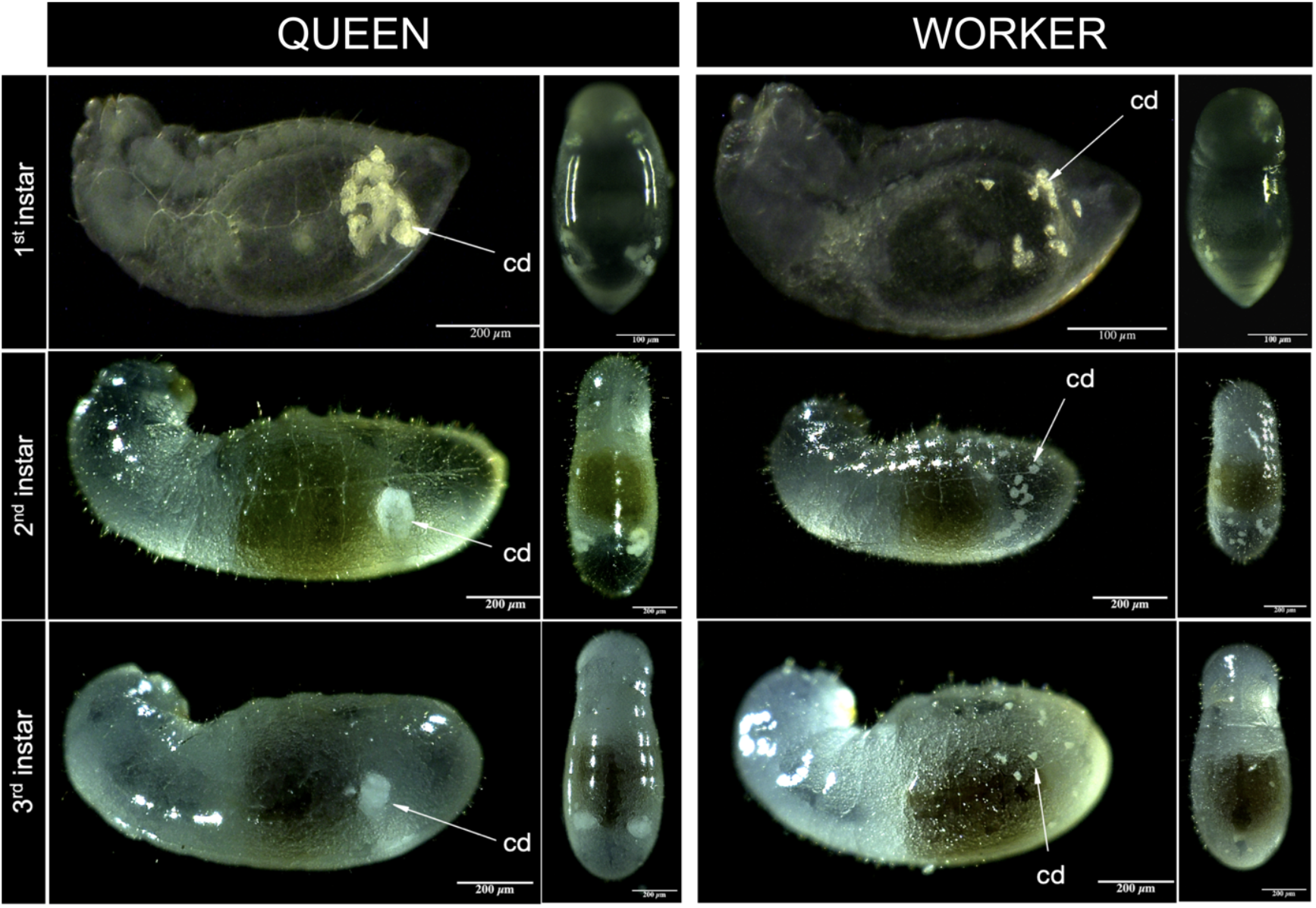
Crystalline deposit patterns distinguish queen- and worker-destined larvae in the ant *Cardiocondyla obscurior*. Stereomicroscope images of queen-destined and worker-destined larval instars show a pair of large, clumped crystalline deposits (cd) in the posterior end of queens, and small, randomly distributed crystalline deposits in workers (lateral and dorsal views).

**Figure 2:**
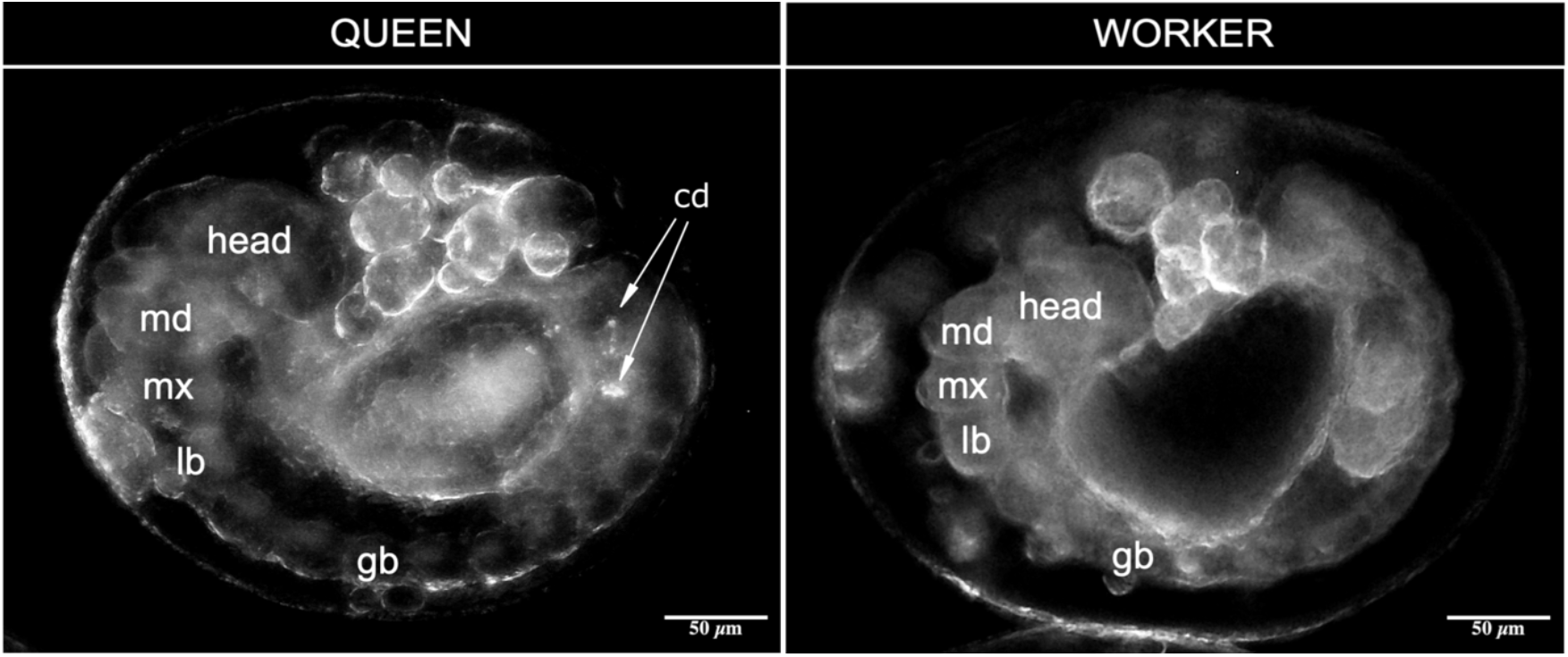
Crystalline deposit patterns in queen- and worker-destined embryos in the ant *Cardiocondyla obscurior*. Queen-destined embryos have crystalline deposits (cd) localized in the last segments of their germ band (gb). Worker-destined embryos do not show any crystalline deposits. The first segments of the embryo are separated into the head and the three gnathal segments: mandibular (md), maxillary (mx) and labial (lb).

**Figure 3:**
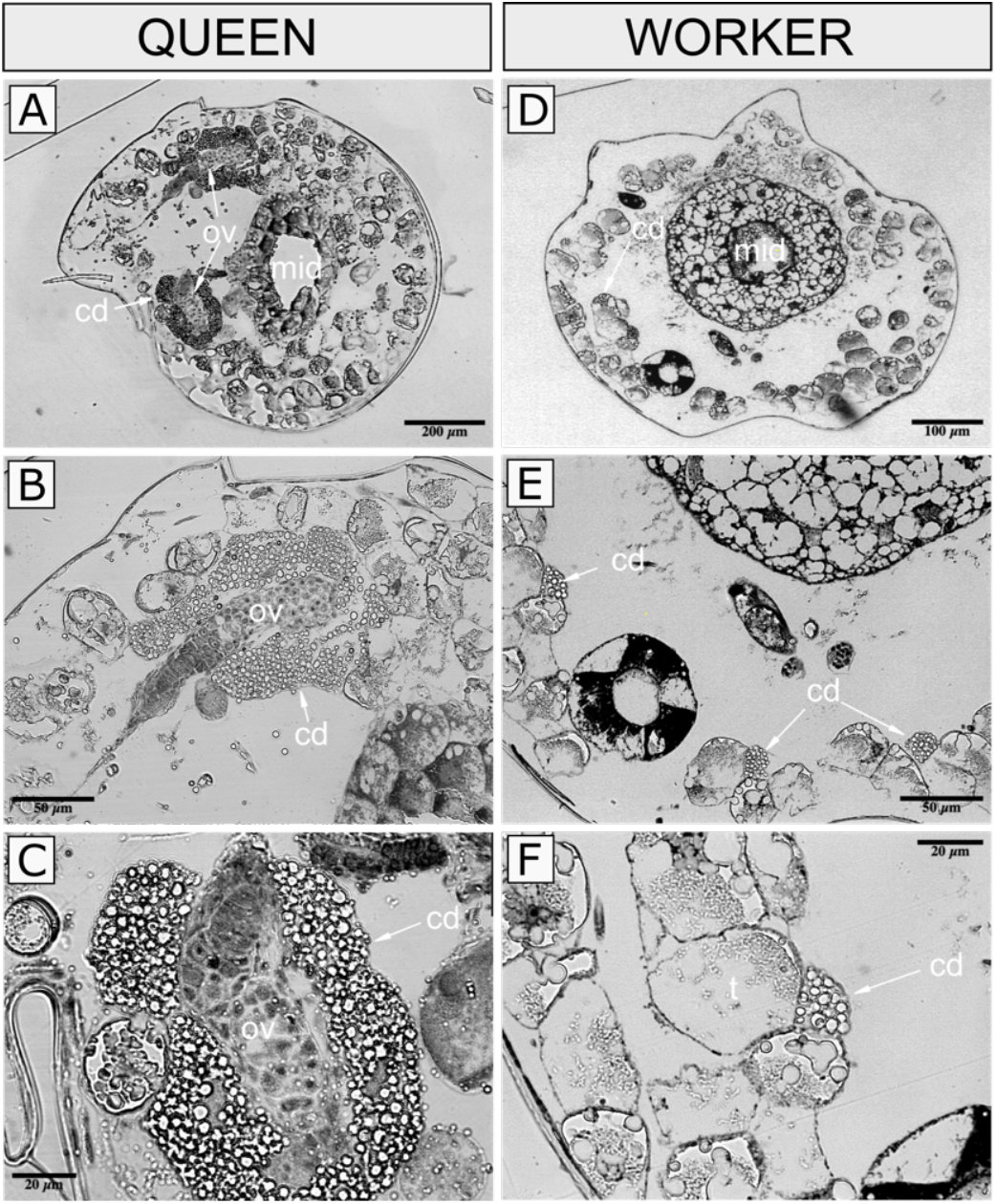
Histological sections of queen and worker-destined third instar larvae. (A) Queen-destined larvae show paired ovaries (ov) close to the midgut (mid). (B,C) Ovaries in queen-destined larvae are surrounded by large crystalline deposits. (D) In workers ovaries are missing. (E, F) In worker-destined larvae crystalline deposits are distributed randomly and aggregate close to trophocytes (t). top row: 5x magnification, middle row: 20x magnification, bottom row: 40x magnification.

Crystalline deposit patterns can be used to identify individuals by caste with high accuracy, and workers successfully reared both castes at comparable rates in colony fragments without queens (Table 1). This demonstrates that caste fate is fixed in the embryo, likely under physiological control of the queen, and provides a first indication that workers do not discriminate between developing queens and workers.

**Table 1:** Localization of crystalline deposits predicts caste in eggs and larvae of the ant *Cardiocondyla obscurior*

| Development Stage | Crystalline deposit pattern | n | Proportion of pupae produced | Survival differences between predicted castes |  | Proportion of correct predictions | Accuracy of caste prediction |  |
| --- | --- | --- | --- | --- | --- | --- | --- | --- |
|  |  |  |  | Odds ratio | p |  | Odds ratio | p |
| egg | queen-like | 100 | 56% (56/100) | 0.664 | 0.157 | 82.1% (46/56) | 83.033 | <0.001 |
|  | worker-like | 100 | 45% (45/100) |  |  | 95.1% (39/41) |  |  |
| 1st instar larva | queen-like | 100 | 47% (47/100) | 0.961 | 1 | 57.4% (27/47) | 28.573 | <0.001 |
|  | worker-like | 100 | 46% (46/100) |  |  | 95.6% (44/46) |  |  |
| 2nd instar larva | queen-like | 170 | 58.2% (99/170) | 0.648 | 0.052 | 96.9% (96/99) | 2133.229 | <0.001 |
|  | worker-like | 175 | 47.4% (83/175) |  |  | 98.8% (82/83) |  |  |
| 3rd instar larva | queen-like | 50 | 60% (30/50) | 0.497 | 0.137 | 100% (30/30) | infinite | <0.001 |
|  | worker-like | 40 | 42.5% (17/40) |  |  | 100% (17/17) |  |  |

Note that 4 out 45 of eggs predicted to be workers developed into males; these were excluded prior to analysis of caste prediction accuracy. No males developed in any of the other setups.

Similar crystalline deposit patterns that reliably predict caste were found in third instar larvae of *C. nuda, C. venustula* as well as *C. wroughtonii*, the sister species to *C. obscurior* (Figure S2, Table S1). No caste-specific patterns were found in *C. “argyrotricha”* (provisional name of a recognized morphospecies to be described by B. Seifert), *C. elegans, C. emeryi, C. minutior* and *C. thoracica* (Figure S3, Table S1), or in any of the other 42 screened ant species (Figure S4, Table S2). This patchy pattern does not follow the phylogenetic relationships within the *Cardiocondyla* genus [33]. Caste-specific patterns are also not linked to the presence of the main bacterial symbiont *Cand*. Westeberhardia cardiocondylae, an endosymbiont with a highly reduced genome that resides in gut-associated bacteriomes, as well as in ovaries in queens [34]. *Cardiocondyla minutior* carries the symbiont [35] but does not display caste-specific patterns, while a *C. obscurior* lineage that is naturally free of *Cand*. Westeberhardia [34,36] and has been maintained in the lab since 2011 shows the patterns (Schultner, pers. obs.). This does not rule out a connection between symbiont but indicates that a potential link must entail more than mere symbiont presence. Future studies will uncover the metabolic processes from which the deposits stem and reveal their function.

### Phenotypes of developing queens and workers

General morphologies of ant larvae have been extensively described, e.g. [37–40], and non-invasive identification of larval sex has been demonstrated previously [41]. Traits that distinguish late-stage worker from sexual larvae (males, queens) have been discovered in six ant species from three subfamilies; these are largely continuous measures, including body size (*Aphaenogaster senilis* [42]; *Harpegnathos saltator* [43]; *Myrmica rubra* [44]), cuticle “shininess” (*Linepithema humile*, [45]), hair density (*Monomorium pharaonis* [46]), and hair morphology *(Acromyrmex echinatior* [47]). Apart from the crystalline deposits, *C. obscurior* queen and worker larvae appeared similar in their morphology, and neither the density nor the morphology of hairs differed between castes in any of the three instars (Figure S5). There was also no difference in size between the castes during early development (Figure 4, eggs: Linear mixed model, length: χ^2^=0.123, df=1, p=0.726; width: χ^2^=0.007, df=1, p=0.934; L1 larvae: ANOVA, head width: F_67,1_=3.009, p=0.088; body length: F_52,1_=0.366, p=0.548). However, as expected in an ant with morphologically distinct queen and worker castes [48], queen larvae grew larger than worker larvae as development progressed (Figure 4, L2 larvae: ANOVA, head width: F_60,1_=3.124, p=0.088; body length: F_60,1_=14.542, p<0.001; L3 larvae: ANOVA, head width: F_60,1_=17.198, p<0.001; body length: F_60,1_=11.754, p<0.01). Third instar queen larvae were also heavier than worker larvae (Figure 5A, L2 larvae: ANOVA, F_10,1_=3.547, p=0.096; L3 larvae: ANOVA, F_10,1_=98.208, p<0.001) but there was no difference in protein content between the two larval castes (Figure 5B, L2 larvae: ANOVA, F_41,1_=0.133, p=0.717; L3 larvae: ANOVA, F_49,1_= 0.876, p=0.354).

**Figure 4:**
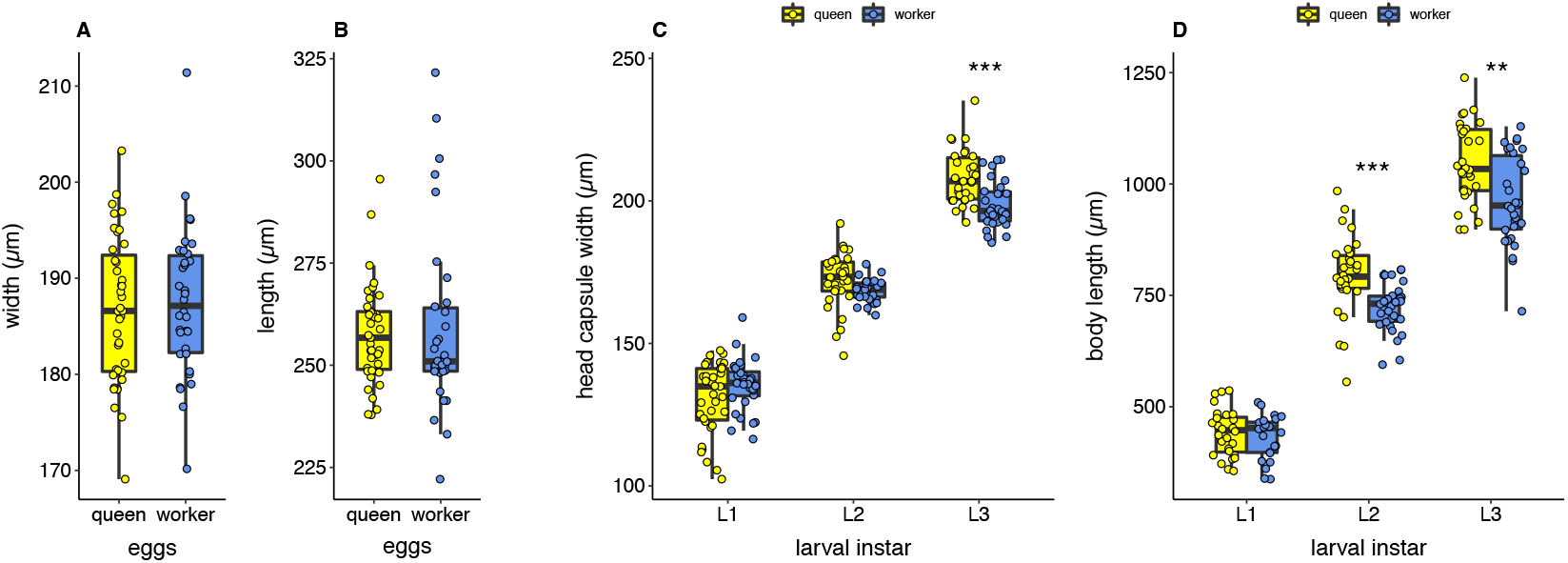
Size of *Cardiocondyla obscurior* queen and worker eggs and larvae (A) egg width, (B) egg length, (C) larva head capsule width, (D) larva body length. L1: first instar, L2: second instar, L3: third instar

**Figure 5:**
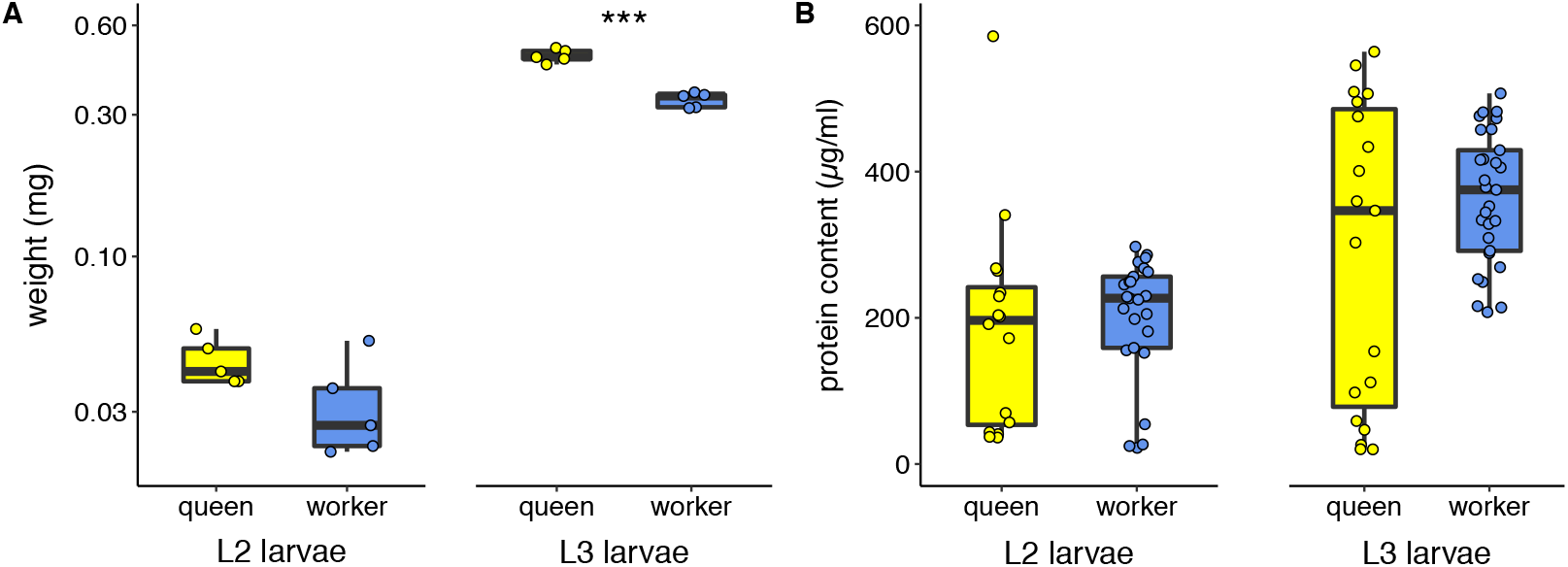
Weight (A) and protein content (B) of queen and worker *Cardiocondyla obscurior* larvae. L2: second instar, L3: third instar

Chemical analyses of cuticular odors of queen and worker third instar larvae and adults of *C. obscurior* detected 32 compounds (Table 2). The profiles of adult queens and workers were slightly more complex than those of larvae (adults: >30 compounds including trace amounts below 0.5%; larvae: 24 compounds). Larval profiles were dominated by n-alkanes, with pentacosane and heptacosane constituting over 65% of the total profile. These two compounds were also present in high relative amounts in adults, together with unsaturated and methyl-branched hydrocarbons. Principal component analysis clearly separated samples by development stage on PC1, apart from two adult samples, which clustered with larva samples for unknown reasons (Figure 6, Table S2). Differences between development stages were confirmed by analysis of similarities (adult vs. larva profiles: ANOSIM, R=0.628, p<0.001). PC2 separated adult castes, with a single worker sample clustering with queens (Figure 6, Table S2, ANOSIM, R=0.548, p<0.001). In contrast, the profiles of queen larvae and worker larvae did not differ (Figure 6, Table S2, ANOSIM, R=-0.059, p=0.651). A similar lack of caste-specific larval odors has been demonstrated in late-stage larvae of the ant *Aphaenogaster senilis* [42], while in *Harpegnathos saltator*, odor profiles of queen-destined larvae were distinct from those of worker-destined larvae, allowing workers to identify larvae by caste [43]. Ant larvae are often considered chemically insignificant because they are easily accepted across colonies, populations and even species (reviewed in [26]. At the same time, brood odors can play an important role in queen fertility signaling within colonies, for example when eggs and larvae carry odors that match those of queens [49,50]. Because of the relative difficulty of distinguishing ant brood by developmental stage, sex and caste, detailed data remain limited, and more studies are needed to understand the function of brood odors.

**Table 2:** Cuticular hydrocarbon profiles of queen and worker larvae and adults of the ant *Cardiocondyla obscurior*

| RI | Compound | mean relative amounts ± standard deviation (%) | | | |
| --- | --- | --- | --- | --- | --- |
|  |  | Queen larvae | Worker larvae | Queens | Workers |
| 2189 | C22 | 3.78 ± 2.66 | 3.89 ± 2.83 | 2.11 ± 1.29 | 6.25 ± 2.49 |
| 2300 | A<sup>a</sup> | 1.2 ± 0.72 | 0.83 ± 0.57 | 0.51 ± 0.12 | 1.49 ± 0.52 |
| 2400 | C24 | 2.97 ± 1.71 | 2.77 ± 1.68 | 1.63 ± 0.89 | 3.28 ± 0.64 |
| 2499 | C25 | 50.49 ± 7.14 | 50.37 ± 4.92 | 26.28 ± 8.93 | 13.48 ± 6.99 |
| 2532 | 13-Me C25<sup>b</sup> | tr | tr | 0.51 ± 0.16 | 1.25 ± 0.64 |
| 2572 | 3-Me C25 | 3.27 ± 1.68 | 2.61 ± 1.14 | 4.66 ± 1.22 | 8.04 ± 1.28 |
| 2599 | C26 | 3.13 ± 0.96 | 3.18 ± 1.19 | 2.47 ± 0.82 | 3.5 ± 0.53 |
| 2632 | B | tr | tr | tr | 0.64 ± 0.41 |
| 2662 | C | tr | tr | tr | tr |
| 2675 | x,x-C27:2 | 1.1 ± 0.42 | 0.59 ± 0.31 | 7.28 ± 1.27 | 13.24 ± 7.64 |
| 2699 | C27 | 17.89 ± 4.13 | 20.8 ± 3.68 | 25.68 ± 5.05 | 9.65 ± 6.68 |
| 2731 | 13-Me, 11-Me C27 | 1.46 ± 0.76 | 0.94 ± 0.44 | 3.3 ± 1.39 | 12.56 ± 5.46 |
| 2749 | 5-Me C27 | tr | tr | tr | 0.58 ± 0.26 |
| 2771 | 3-Me C27 | 1.79 ± 0.37 | 1.61 ± 0.35 | 2.66 ± 0.5 | 2.35 ± 0.42 |
| 2799 | D<sup>a</sup> | 1.63 ± 0.74 | 1.5 ± 0.75 | 1.89 ± 0.73 | 2.69 ± 0.8 |
| 2831 | E | tr | tr | tr | 1.06 ± 0.51 |
| 2861 | 4-Me C28 | 0.61 ± 0.27 | 0.6 ± 0.44 | tr | tr |
| 2880 | x-C29:1 | tr | tr | tr | tr |
| 2899 | C29 | 3.7 ± 1.33 | 4.3 ± 1.07 | 8.73 ± 2.52 | 4.24 ± 2.58 |
| 2929 | 13-Me, 11-Me C29 | 1.02 ± 0.53 | tr | 1.45 ± 0.42 | 5.39 ± 2.21 |
| 2972 | 3-Me C29 | tr | tr | 0.8 ± 0.31 | tr |
| 2999 | C30 | 1.68 ± 0.81 | 1.94 ± 0.8 | 1.61 ± 1.38 | 4.36 ± 1.77 |
| 3006 | F | tr | tr | 0.59 ± 0.19 | tr |
| 3028 | G | tr | tr | tr | tr |
| 3060 | H | tr | tr | tr | tr |
| 3079 | x-C31:1 | tr | tr | tr | tr |
| 3098 | C31 | 0.57 ± 0.18 | 0.54 ± 0.23 | 1.88 ± 0.76 | tr |
| 3125 | 13-Me C31<sup>b</sup> | 0.91 ± 0.42 | tr | 2.28 ± 0.71 | 1.85 ± 0.9 |
| 3172 | 3-Me C31 | tr | tr | tr | tr |
| 3196 | I | tr | tr | 0.66 ± 0.28 | 0.52 ± 0.25 |
| 3297 | C33 | 0.69 ± 0.42 | 0.77 ± 0.61 | 0.93 ± 0.52 | 1.16 ± 0.72 |
| 3326 | 13-Me C33<sup>b</sup> | tr | tr | tr | 0.77 ± 0.42 |

RI - linear retention index
tr – trace amounts (<0.5% of total profile)
<sup>a</sup> diagnostic ions were not detectable, but RIs indicate: A=C23, D=C28
<sup>b</sup> peaks may include a further monomethyl alkane (11-Me) of the respective chain length at low relative concentration

**Figure 6:**
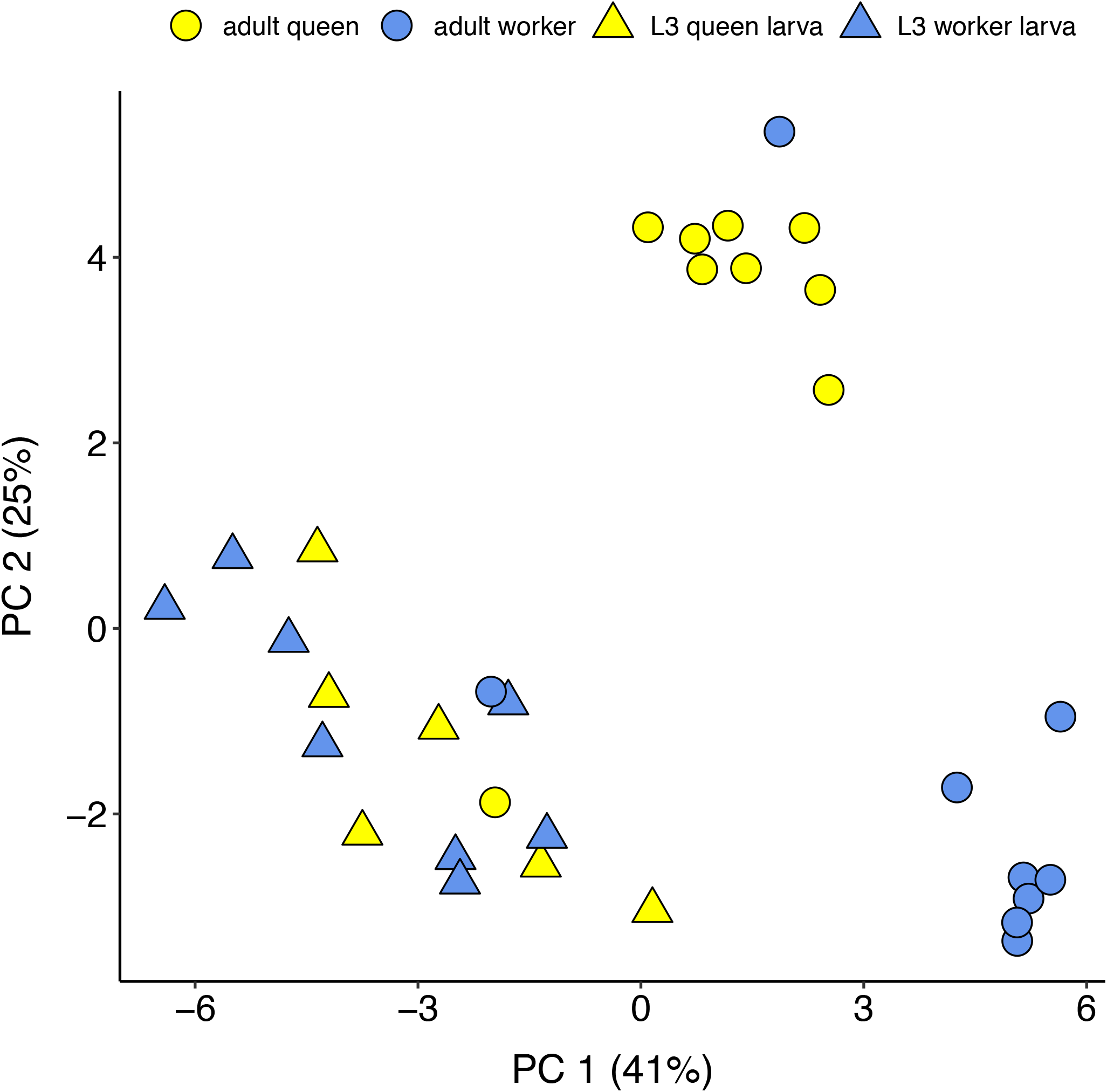
Principal component analysis of cuticular hydrocarbon profiles of queen and worker larvae and adults in the ant *Cardiocondyla obscurior*

### No sign of caste conflict involving brood

In line with the absence of caste-specific morphological and chemical cues, adult workers did not adjust their behavior to the caste of encountered larvae in any of the five tested scenarios. Workers were similarly attracted to queen and worker larvae (Figure S6, number of contacts: ANOVA, F_44,1_=3.193, p=0.077; duration of contacts: Mann-Whitney-U-test, W=1184, p=0.072) and did not preferentially retrieve larvae of either caste (queen larvae retrieved: 59% (40/67), worker larvae retrieved: 40% (27/67), Chi-squared test, χ^2^=2.522, df=1, p=0.113). Observations of feeding behavior revealed no preference for either caste, with 65% (13/20) of worker larvae and 59% (10/17) of queen larvae containing fluorescent dye in their gut two hours after starved colonies composed of larvae, workers and queens were fed with fluorescent honey (Fisher’s exact test, odds ratio=0.774, p=0.745). There was no difference in the overall survival of queen and worker larvae in these setups (survival queen larvae: 68% (17/25); survival worker larvae: 80% (20/25); Fisher’s exact test, odds ratio=0.538, p=0.52). Workers also reared second instar larvae of both castes to pupation with similar success when they were simultaneously presented with queen and worker larvae in the presence of a mated queen (larvae reared to pupation: queens: 38% (19/50); workers: 29% (14/48); Fisher’s exact test, odds ratio=1.482, p=0.398). Finally, when workers reared groups of first and second instar larvae of unknown caste alone or in the presence of a fertile queen, they reared similar proportions of new queens (L1 larvae - queen present: 42.5% queens (17/40); queen absent: 32.6% queens (16/49); Fisher’s exact test, odds ratio=1.517, p=0.382; L2 larvae -queen present: 13.7% queens (10/73); queen absent: 12.7% queens (7/55); Fisher’s exact test, odds ratio=1.089, p=1). Together, these results indicate that *C. obscurior* workers are indifferent to the caste of developing individuals.

## Conclusion

Previous studies in *C. obscurior* found that queens control egg sex ratios, and that workers do not manipulate secondary sex ratios of brood [27,28]. In the present study we show that female caste in this species is determined in the egg stage, indicating that caste ratios are also under maternal control. Developing queens and workers differed in size and weight in late larval stages, but not in cuticular odors, hair morphology or physiology. Workers indiscriminately reared eggs and larvae of both castes in different social contexts, including when a fertile queen was present. *C. obscurior* queens thus appear to have control over sex and caste allocation, two colony-level traits that often elicit conflict in social Hymenoptera.

Queen control of sex allocation has been demonstrated previously in ants [19,51–53], but complete control over caste allocation has, to the best of our knowledge, not been described. In other ant genera with fully sterile workers such as fire ants (*Solenopsis*), Pharaoh ants (*Monomorium*) and big-headed ants (*Pheidole*), caste may similarly be determined in the egg, allowing queens control over primary caste ratios [13,54]; however, workers continue to play an important role in regulating secondary caste ratios by culling sexual larvae in the presence of fertile queens or their pheromonal equivalent [46,55–61].

Why then is *Cardiocondyla* different? Colonies are highly inbred due to the evolution of wingless males with lifelong spermatogenesis, which remain in the nest over the course of their relatively long lives to mate with multiple related queens, including sisters and mothers [62–64]. Inbreeding results in particularly high levels of relatedness, and consequently decreased conflict potential within colonies. *C. obscurior* moreover has a relatively low degree of dimorphism between sexuals and workers [48,65] and queens only have six ovarioles, producing about 1-2 eggs per day and an average of 350 eggs over the course of their lives [29]. Accordingly, colonies are small and contain on average one to several queens and ∼30 workers [65]). In comparison, *Monomorium, Solenopsis* and *Pheidole* queens are much larger than their workers and highly fertile, with ovariole numbers ranging from 24 to over 100 [15,66], and they can produce hundreds of eggs per day, and hundreds of thousands of eggs over the course of their lives. High queen fertility may allow these species to adopt a strategy of brood culling to dynamically regulate sex and caste allocation, possibly under constraints imposed by high dimorphism between sexuals and workers such as the inability to rear sexuals when colonies are small, lack development stages involved in nutrition processing, or resources are limited [56,67,68]. In *C. obscurior*, low queen fertility means individuals eggs have high relative value for all colony members, likely preventing the evolution of a similar strategy. However, low dimorphism between sexuals and workers allows both castes to be produced at comparable costs, so that even tiny colony fragments consisting of only a few workers and brood can initiate a new colony. This provides an important advantage in the exposed and often disturbed arboreal habitats where the ant thrives, and may help explain its success as a global tramp species [69]. How the unique combination of high relatedness, worker sterility, limited queen fertility and ecological constraints resulted in the evolution of maternal control of egg caste, and the physiological mechanisms underlying this process, remain to be uncovered.

## Methods

### Ants

*Cardiocondyla obscurior* and several other *Cardiocondyla* species are cosmotropical tramp ants [70]. This myrmicine genus has evolved wingless fighter males with lifelong spermatogenesis, which remain in the nest and mate with newly-hatched nestmate queens [62]. *Cardiocondyla obscurior* has the smallest known ant genome (∼193MB, [65,71]). Adult queens and workers differ in size and morphology and workers lack ovaries [48]. Larvae develop via three instars, which can be distinguished by body shape and the degree of melanization of the mandibles [72]. All ants used for this study were from the Old World lineage [71,73], maintained in the lab since 2010. Stock and experimental colonies were kept in plaster-bottom nests with nest inserts made of acrylic glass covered by dark foil in a climate chamber under a 12h/12h and 22°C/26°C night/day cycle at 70% humidity. All colonies were provided with water and fed three times a week with honey, pieces of cockroaches and fruit flies.

### Crystalline deposit patterns in queen and worker larvae of *Cardiocondyla*

Localization of crystalline deposits was characterized in eggs, first instar (L1), second instar (L2) and third instar (L3) larvae as queen-like or worker-like by visually inspecting brood from stock colonies under a stereomicroscope. For better detection of the patterns, eggs and L1 larvae were submerged in a dissection dish containing 0.3% PBT, after which they were mounted on a microscope slide and sealed with nail polish. From each developmental stage, representative individuals with queen-like and worker-like patterns were selected and photographed using a stereomicroscope connected to a camera (Keyence VHX 500FD, Neu-Isenburg, Germany). Crystalline deposit patterns were characterized in the same manner in L3 larvae from eight additional *Cardiocondyla* species (*C. “argyrotricha”, C. elegans, C. emeryi, C. minutior, C. nuda, C. thoracica, C. venustula, C. wroughtonii*) spanning the phylogeny [33]. We further examined larvae of six species from four subfamilies available as live colonies and accessed photos of 36 species from seven subfamilies from Alex Wild’s library (https://www.alexanderwild.com/Ants/Natural-History/Metamorphosis-Ant-Brood/), for a total of 51 examined species (Table S2).

In *C. obscurior*, the development of all stages was tracked to confirm that crystalline deposit patterns are associated with caste (Table 1). Eggs and L1 larvae were transferred to filter paper to remove excess buffer before being moved in groups of 10 to experimental colonies containing 10 workers each (eggs: queen-like=100, worker-like=100, L1 larvae: queen-like=100, worker-like =100). L2 and L3 larvae were transferred in groups of 10-15 to experimental colonies containing 10-12 workers (L2 larvae: queen-like=170, worker-like=175; L3 larvae: queen-like=50, worker-like=40). We also tracked the development of L3 larvae from some of the other *Cardiocondyla* species (*C. minutior, C. nuda, C. thoracica, C. venustula, C. wroughtonii*) (Table S1). Experimental colonies were monitored three times per week and, when necessary, workers were added from the corresponding stock colonies to standardize worker number. Developing individuals were monitored until pupation, at which time they were counted and classified according to female caste, until no more brood remained. From these data, rates of survival until pupation and caste ratios were calculated. For *C. obscurior*, caste-specific survival and accuracy of caste prediction were tested with Fisher’s exact tests (function fisher.test in R 4.0.3 [74]).

### Larval histology

*C. obscurior* L3 larvae were collected from stock colonies and sorted according to crystalline deposit patterns. Sorted larvae were transferred into a fixation solution consisting of 25% glutaraldehyde (GAH) in cacodylate buffer [(50 mM cacodylic acid, pH 7.3) containing 150 mM Sucrose] (GAH : cacodylate buffer = 1 : 12,5) and kept overnight at 11 °C. Samples were then rinsed in cacodylate buffer on ice, and fixated in 4% osmium tetroxide in cacodylate buffer. After fixation, larvae were washed in cacodylate buffer, dehydrated in a graded ethanol series and embedded in Epon. Transversal semithin sections of 1 µm were cut and stained with methylene blue and Azur II (Richardson’s stain). Semi-thin sections were scanned with a Zeiss Primo Star microscope and imaged with a Moticam 580 digital microscope camera.

### Phenotypes of *C. obscurior* queen and worker-destined larvae

#### Size

Length and width of eggs with queen-like and worker-like crystalline deposit patterns were assessed (n=39 queen-like eggs from 7 stock colonies; n=34 worker-like eggs from 7 stock colonies). Each egg was photographed through a stereomicroscope connected to a camera at 200x magnification and its length and width measured using the system software (Keyence VHX 500FD, Neu-Isenburg, Germany). Caste-specific egg lengths and widths were compared using linear mixed models with length or width as response variables and larval caste as explanatory variable. To account for between-colony variation, the stock colonies from which eggs originated were implemented as random variables. Model residuals were checked for normality using Shapiro-Wilks tests (function shapiro.test) and residual tests implemented in the DHarma package (function testResiduals [75]) in R 4.0.3 [74]. Size differences between castes were assessed using a Wald **Χ**^2^ test (function Anova, package car [76]).

To assess the size of queens and workers over the course of larval development, head capsule width and body length were measured (L1 larvae: n=35 queens, n=32 workers; L2 larvae: n=30 queens, n=30 workers; L3 larvae: n=30 queens, n=30 workers). Individuals were collected from stock colonies and sorted by instar and caste, and then anaesthetized with CO_2_ for four hours. Larvae were placed on their backs and photographed using a JEOL scanning electron microscope (L1 larvae) or a stereomicroscope connected to a camera (L2 & L3 larvae). Measurements were taken using ImageJ [77] after setting the appropriate scales. In each instar, head capsule width and body length of queen and worker larvae were compared using linear regressions with width or length as response variables and larval caste as explanatory variable. Model residuals were checked for normality using Shapiro-Wilks tests (function shapiro.test) and residual tests (function testResiduals [75]) in R 4.0.3 [74]. Size differences between castes were assessed using a Wald **Χ**^2^ test (function Anova, package car [76]).

#### Weight

Five pools of five larvae each per instar (L2, L3) and caste (queen, worker) were collected from stock colonies and weighed on a Sartorius SC2 fine scale (Sartorius, Göttingen, Germany). Weight differences were analysed within each instar with a linear regression using weight per individual larva in mg as response variable and caste as explanatory variable. Model residuals were checked for normality using the Shapiro-Wilks test (function shapiro.test) and residual tests (function testResiduals [75]) and differences between castes were assessed using a Wald **Χ**^2^ test (function Anova, package car [76]) in R 4.0.3 [74].

#### Hair density and morphology

Hair density and morphology was assessed qualitatively in queen and worker larvae of all three instars (n=3 per caste and instar). Larvae were collected from stock colonies and sorted by instar and caste, and then placed on a self-adhesive sample insert and moved to -20°C for seven minutes. Freshly killed larvae were scanned with a JEOL scanning electron microscope and density and morphology of hairs was compared visually.

#### Protein content

L2 and L3 queen and worker larvae were collected from stock colonies and fixed individually in 15 µl of 1x PBS solution and frozen at -20°C until further analysis (n=30 per instar and caste). For analysis the samples were allowed to thaw on ice and were then homogenized using pestles. Protein concentration in the samples was measured using the Bicinchoninic Acid Protein Assay Kit (Sigma Aldrich, St. Louis, MO, USA) according to the manufacturer’s protocol with the following modifications: half of the volume for the microplate assay was used, resulting in 100 µl of working reagent and 12.5 µl of sample consisting of 3 µl of homogenized ant sample and 9.5 µl of 1x PBS. All samples were run in two technical replicates at an absorbance of 526 nm using a SpectraMax iD3 microplate reader (Molecular Devices, San Diego, CA, USA). The average of the two technical replicates was used for further calculations. Protein concentration was calculated using a standard curve created with BSA (bovine serum albumin). Within each instar, protein content of queen and worker larvae was compared using a linear regression with protein content in µg/ml as response variable and larval caste as explanatory variable. Model residuals were checked for normality using a Shapiro-Wilks test and residual tests implemented in the DHARMa package. Differences between castes and instars were assessed with a Wald **Χ**^2^ test. A Levene’s test was used to compare the variances of protein content between castes; this was done separately for each instar.

#### Cuticular odors

Cuticular hydrocarbon (CHC) profiles of pools of L3 queen-destined larvae (20 individuals each, n=6), L3 worker-destined larvae (20 individuals each, n=8), adult queens and adult workers (10 individuals each, n=9 for both groups) were analyzed using gas chromatography coupled with mass-spectrometry (GC-MS). For each sample, individuals from at least five stock colonies were pooled to account for colony-level differences. To obtain samples free of contamination from nest material, individuals used in analyses were collected from stock colonies, separated by caste and kept in freshly-plastered experimental nests for ∼24h prior to sampling. All samples were frozen in glass screw top vials and stored at -20°C until extraction.

Samples were extracted in 100 µl HPLC grade hexane for 1 minute. Hexane extracts were then fractionated using silica columns (Chromabond SiOH, 1 mL/100mg). Columns were rinsed consecutively with 1 ml of hexane, dichloromethane, methanol and again dichloromethane, before being conditioned with 2 ml of hexane and adding the sample. The non-polar fraction containing hydrocarbons was eluted with 1 ml of hexane, and subsequently evaporated under a stream of nitrogen. After being re-dissolved in 100 µl of hexane, 1 µl per sample was subjected to GCMS (Shimadzu GCMS-QP2010 Plus) on a SGE BPX-5 column (30m × 0.25µm × 0.25mm). Helium gas (1 mL/min) served as carrier gas, and injection was splitless. The temperature program started at 70 °C isothermal for 1 min, after which the temperature was raised to 200 °C at 30 °C/min and then from 200 to 320 °C at 5 °C/min, where it was held for 5 min.

Compound determination of peaks was based on their characteristic mass spectra and matching of retention indices (for methyl-branched alkanes: [78]. For peaks of a concentration too low to yield an unambiguous mass spectrum, compounds were considered to be identical to those present in other samples if their retention indices matched. Compounds that could not be identified were labeled with letters, while double-bond positions of unsaturated hydrocarbons were not further elucidated and are indicated with “x” (Table 2). Only peaks with retention times over 15 minutes (docosane and longer chain lengths) were used for quantitative analyses. Relative peak areas were calculated by dividing individual peak areas by the summed area of all hydrocarbon peaks. Values were then square-root transformed, centered and scaled prior to principal component analysis (PCA, function prcomp in R). Differentiation between groups was tested on square-root transformed values with analysis of similarities (ANOSIM, function anosim in R); tests were run separately for comparisons between developmental stages (adults vs. larvae) and castes within each stage (adult queens vs. adult workers, queen larvae vs. worker larvae).

### Worker behavior toward queen and worker larvae in different in social contexts

The behavior of adults toward queen and worker larvae was investigated in five scenarios.

First, we investigated whether isolated worker ants are differentially attracted to L3 queen and worker larvae. This was done in round observation chambers (diameter 20 mm), which were visually divided into two equally sized halves. In the center of each half, a clean piece of plastic foil with either a L3 queen or worker larva from the same stock colony was placed, so that each chamber contained one larva of each caste. Larvae were glued to the plastic foil with a small drop of sucrose solution to prevent workers from moving larvae around during the observation period. To avoid potential side bias, queen and worker larvae were randomly placed in the left and right halves of the chambers. Once larvae were placed in the chamber, one worker from the same stock colony was added and a transparent microscope slide placed on top to prevent workers from escaping. Chambers were observed for 12 minutes and the number and duration of contacts, i.e., instances of bodily contact between the adult worker and either queen or worker larva, were scored using JWatcher software [79] (n=44 chambers observed in total). The number of contacts was analysed with a linear regression using larval caste as the explanatory variable. Model residuals were checked for normality using the Shapiro-Wilks test and residual tests implemented in the DHARMa package. The duration of contacts did not follow a normal distribution and was analysed with a Mann-Whitney-U-test (function wilcox.test) in R.

Second, we tested whether worker ants retrieve L3 queen and worker larvae at different rates. From each of five stock colonies, one experimental colony was set up containing 12 workers and one mated queen. Colonies were allowed to settle for 48h. One L3 worker larva and one L3 queen larva from the original stock colonies were then added to the corresponding experimental colonies on a clean piece of filter paper. Experimental colonies were observed until the first larva was retrieved into the nest and the caste of the retrieved larva was noted. This procedure was repeated at least 11 times in each experimental colony, until a total of 67 retrieval events were documented. Retrieval rates of the two castes were compared using a **Χ**^2^ Test.

Third, we investigated whether workers display preferential treatment towards larvae of either caste under food limitation. Five experimental colonies were set up, each containing one mated queen, five workers, and five L2 larvae of each caste. Colonies were then starved of sucrose for three days. On day three, colonies were presented with 0.25 µl of a 1.5 M sucrose solution containing 0.08 g l^-1^ Rhodamine B (Czaczkes et al., 2019). After two hours, all larvae were counted and examined for the presence of Rhodamine B in the digestive tract under a green laser light (555 nm) using a Zeiss Axioplan-2 fluorescence microscope. Larvae positive for Rhodamine B were considered “fed”, while the absence of fluorescence in larvae was scored as “unfed”. The number of fed and unfed larvae of each caste, as well as the overall survival of queen and worker larvae, was compared using Fisher’s exact tests (function fisher.test in R).

Fourth, we assessed whether workers changed their rearing behavior in the presence of a mated queen when presented with queen and worker larvae simultaneously. Five colonies were set up, each with 10 second instar queen and 10 second instar worker larvae, as well as 20 workers and one mated queen from the same stock colony. Colonies were fed as described above and all pupae counted and their sex and caste noted. Caste-specific survival of larvae was compared using Fisher’s exact test (function fisher.test in R).

Fifth, we tested whether the presence of a mated queen influenced secondary caste ratios in brood of unknown caste. First and second instar larvae of unknown caste were collected from stock colonies and reared in experimental colonies with and without queens. For each instar separately, colonies were set up containing 15 larvae and 15 workers, as well as one mated queen (L1 n=10, L2 n=8) or no queen (L1 n=13, L2 n=9). In each experimental colony, all individuals came from the same stock colony. Colonies were fed as described above and all pupae counted and their sex and caste noted. The proportion of queens produced in colonies with and without a mated queen was compared using Fisher’s exact test (function fisher.test in R).

## Acknowledgements

We thank B. Lautenschläger and M. Schiwek for help with histology, E. Strohm for help identifying the crystalline nature of the deposits, and A. Koch, K. Pogorelski, A. Reisinger and C. Winkler for help with experiments. JO dedicates this paper to Margit Oettler, for nature and nurture. This study was funded by DFG grants (DFG OE 549/3-1, OE 549/3-2, OE 549/3-3).

## Figure legends

**Supplementary Figure S1:**
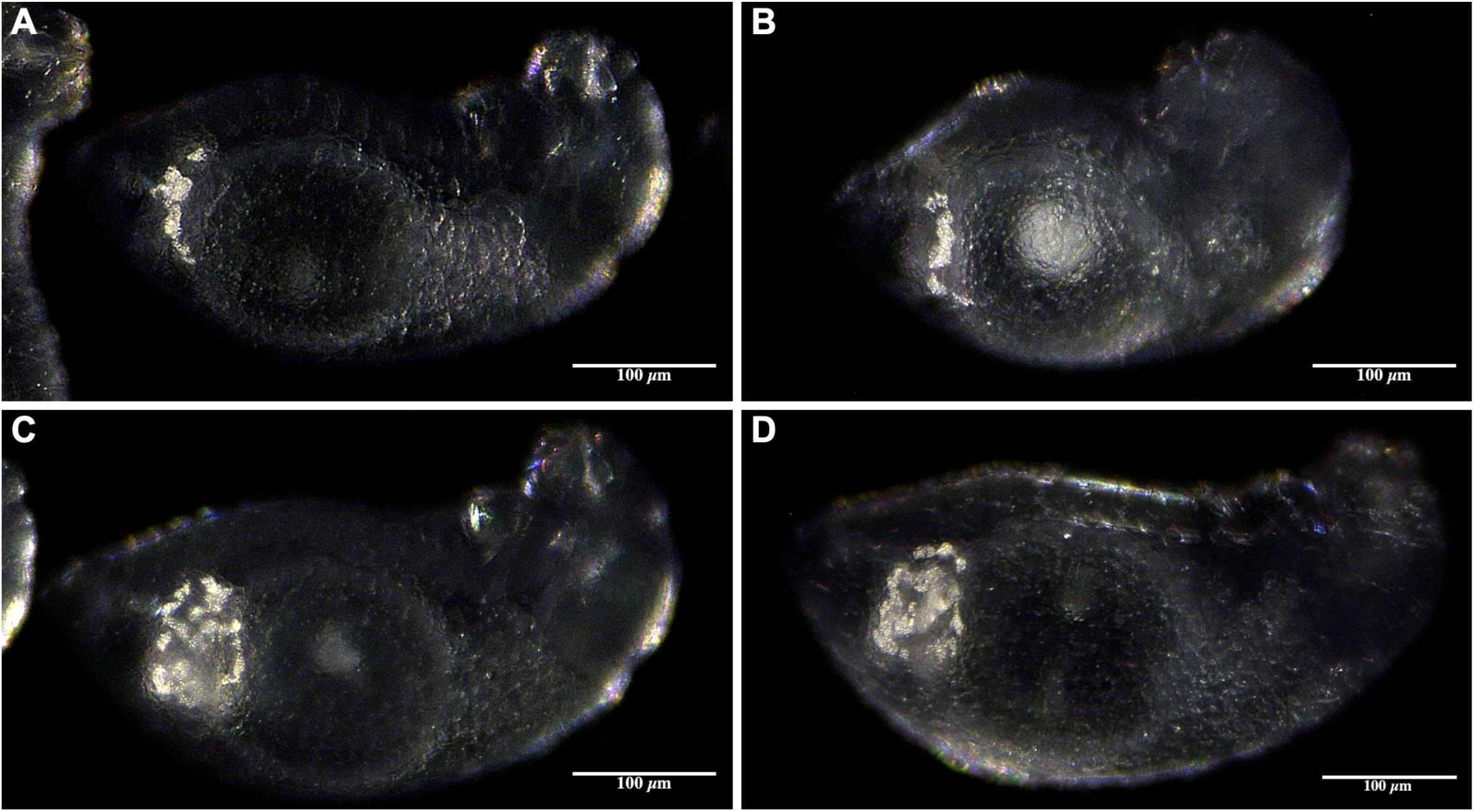
Crystalline deposit patterns are variable in first instar queen-destined larvae. (A,B) String of pearl-like deposits. (C,D) Blob-like deposits.

**Supplementary Figure S2:**
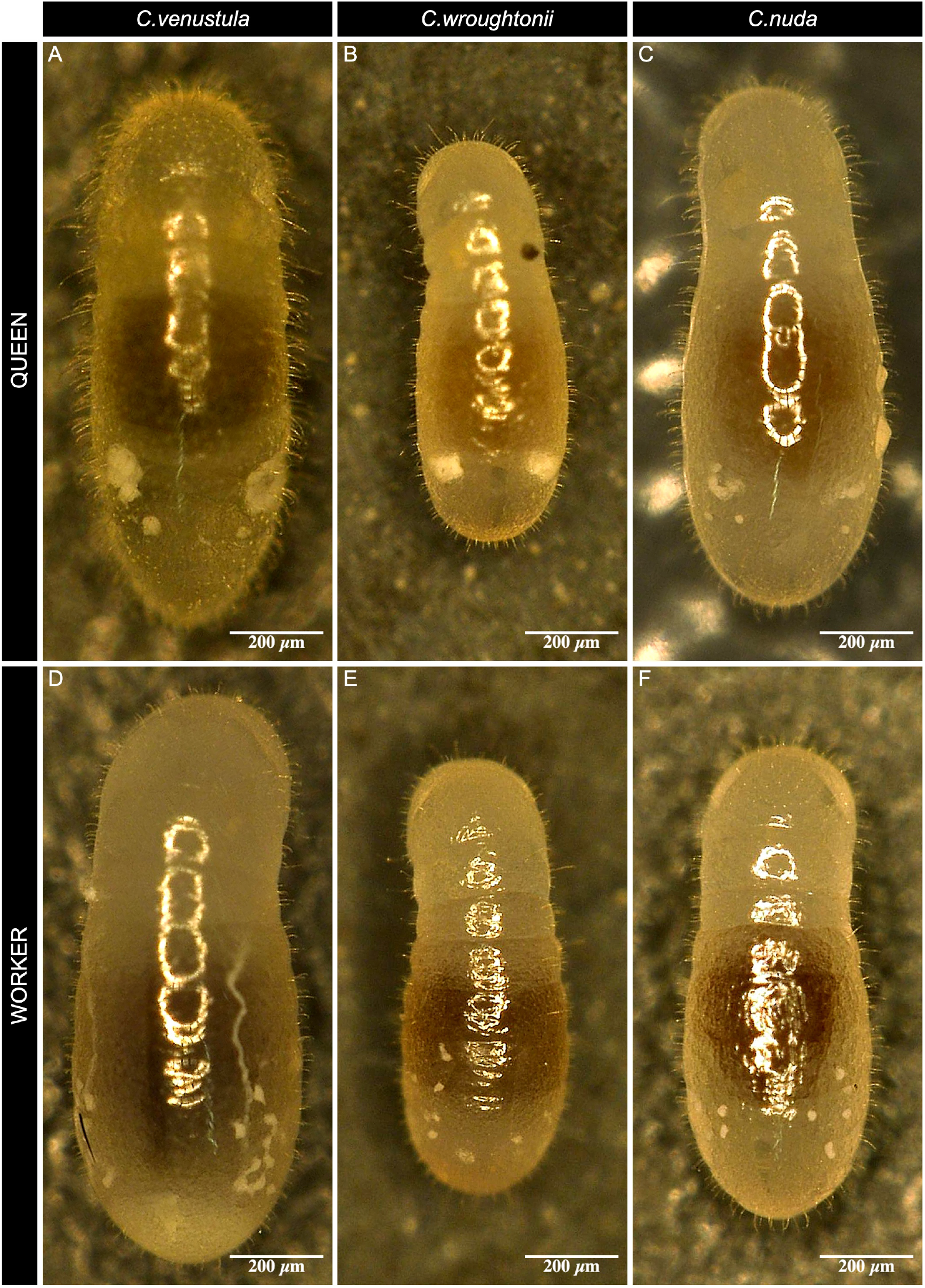
Crystalline deposit patterns distinguish queen- and worker-destined larvae in several *Cardiocondyla* species. Stereomicroscope images of queen-destined (A, B, C) and worker-destined (D, E, F) third instar larvae in *C. venustula, C. wroughtonii* and *C. nuda*.

**Supplementary Figure S3:**
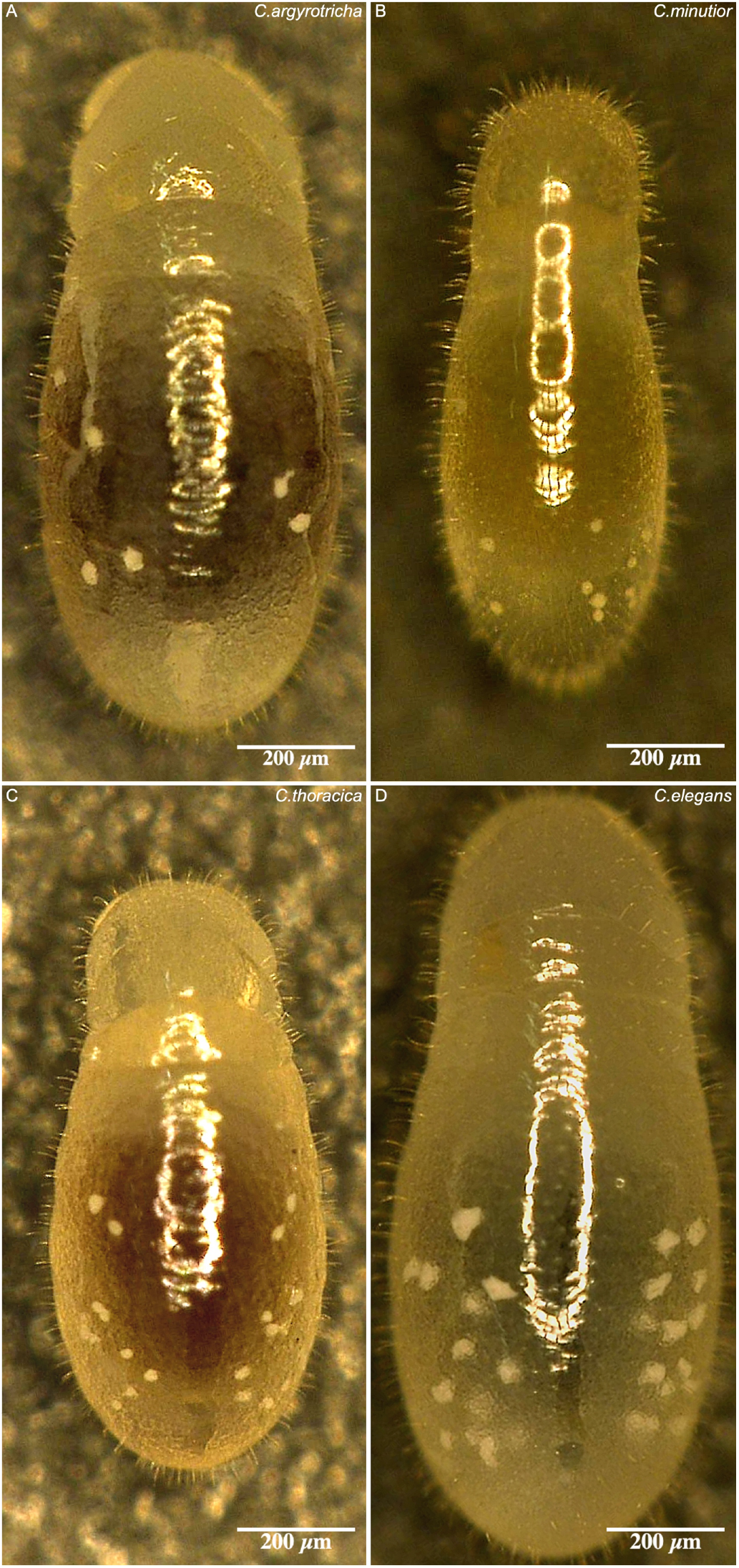
Crystalline deposit patterns in *Cardiocondyla* species without caste specific patterns. Stereomicroscope images of third instar larvae in *C. “argyrotricha”, C. minutior, C. thoracica* and *C. elegans*. No pictures were taken of *C. emeryi*.

**Supplementary Figure S4:**
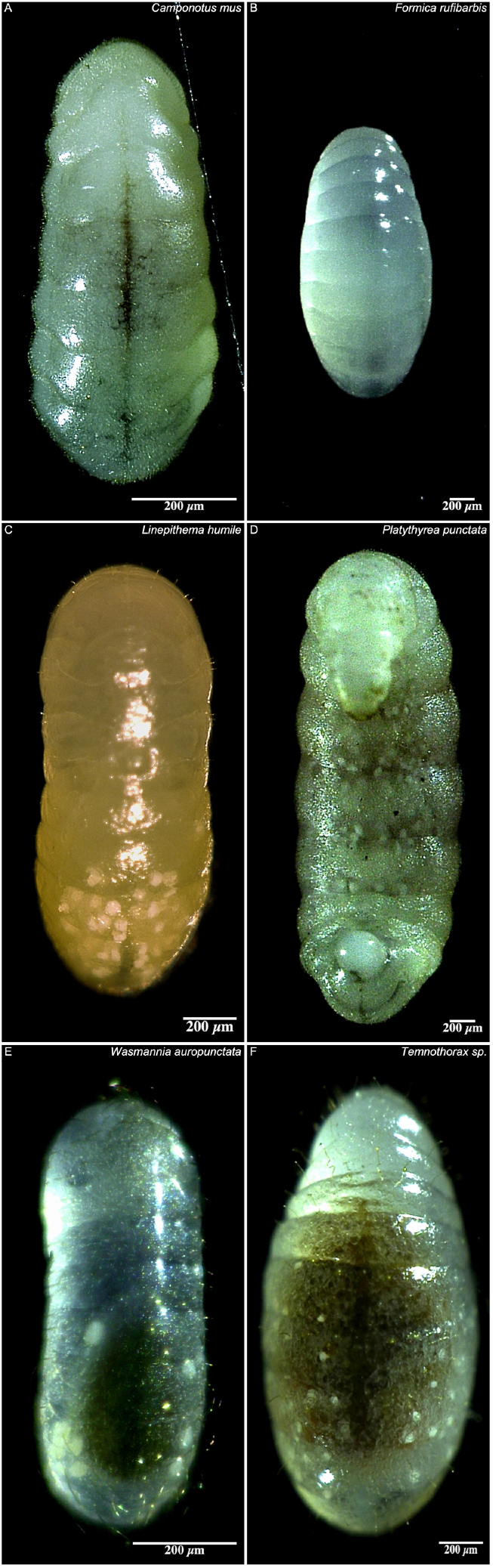
Crystalline deposit patterns in seven species from four ant subfamilies available as live colonies in the lab. Stereomicroscope images of larvae in *Camponotus mus* (Formicinae), *Formica rufibarbis* (Formicinae), *Linepithema humile* (Dolichoderinae), *Platythyrea punctata* (Ponerinae) (ventral view), *Wasmannia auropunctata* (Myrmicinae), *Temnothorax sp*. (Myrmicinae). Both representatives of the Formicinae lack visible deposits.

**Supplementary Figure S5:**
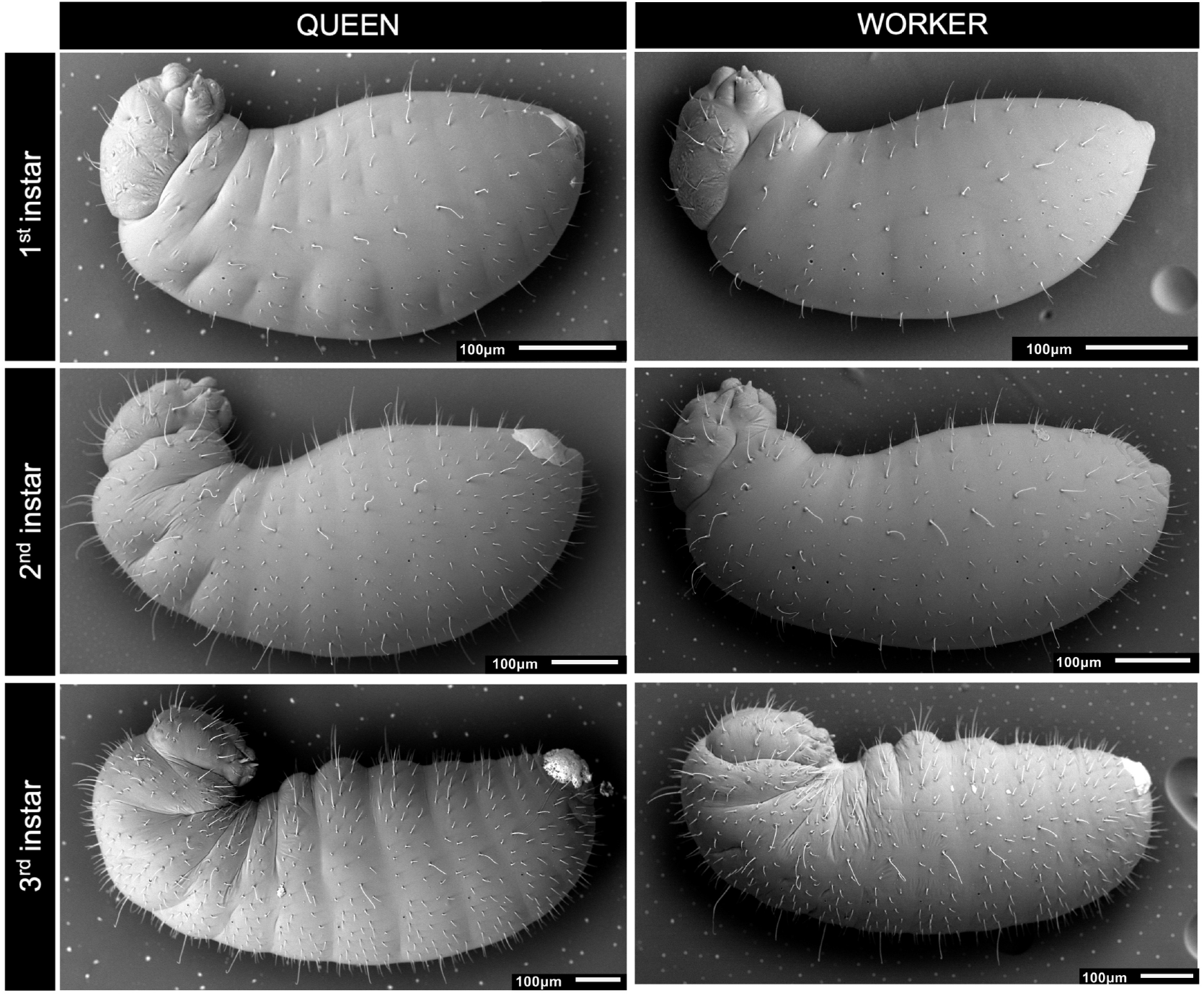
Transmission electron microscopy reveals similar external morphologies in queen and worker destined larvae of the ant *Cardiocondyla obscurior*

**Supplementary Figure S6:**
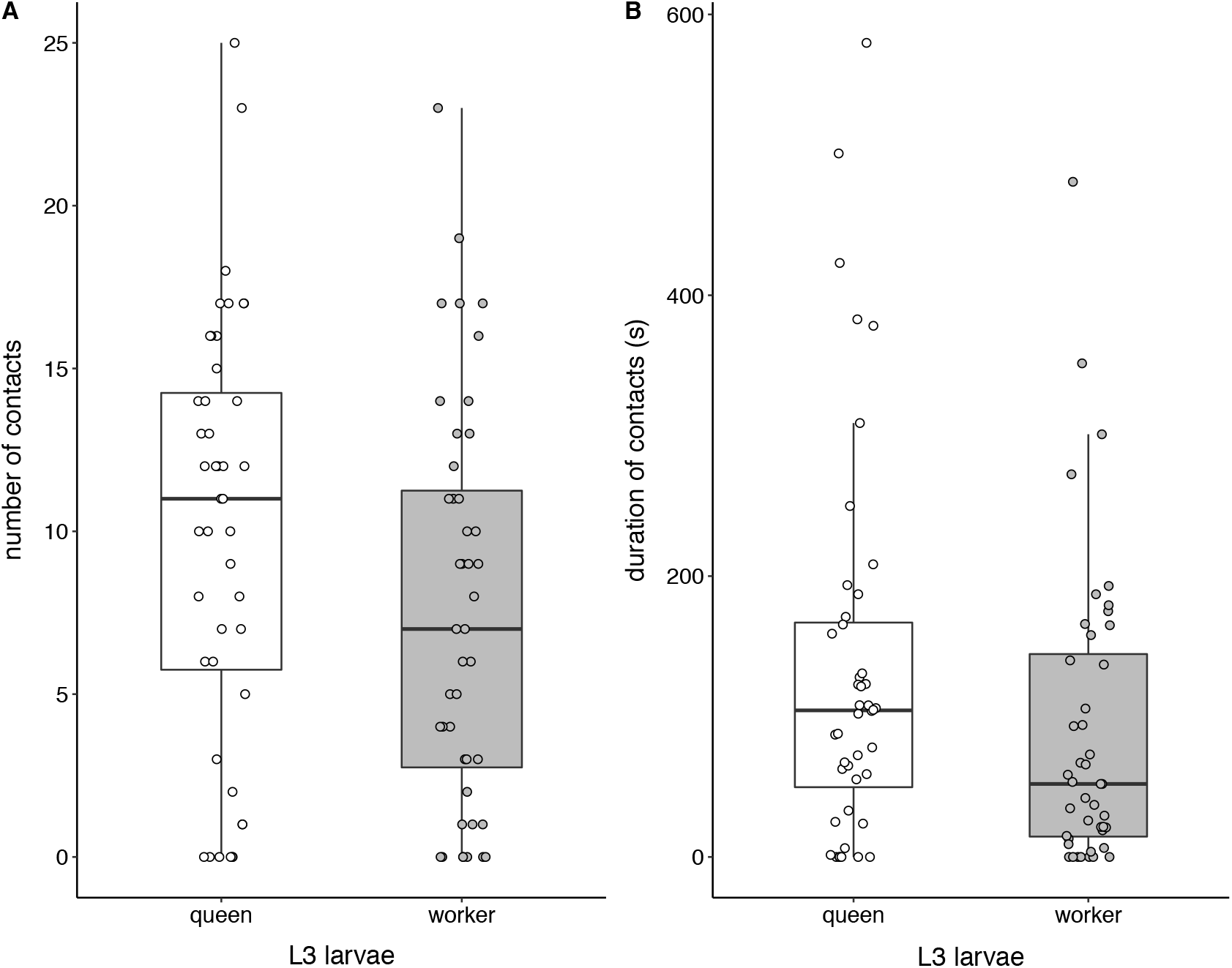
Workers are similarly attracted to queen and worker third instar larvae. (A) number of contacts, (B) duration of contacts

**Supplementary Table S1:**
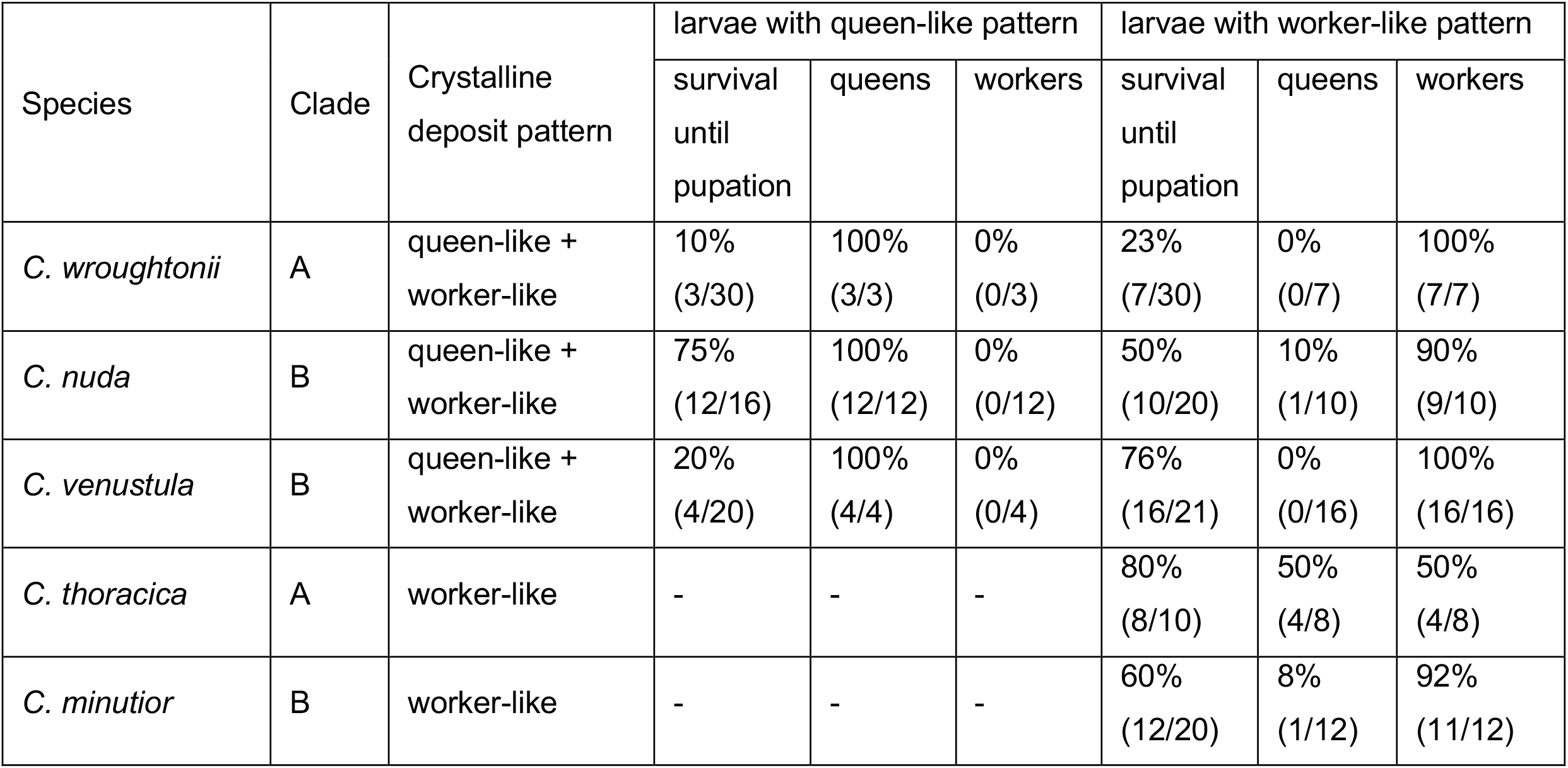
Crystalline deposit patterns in third instar larvae predict caste in some *Cardiocondyla* species

**Supplementary Table S2:**
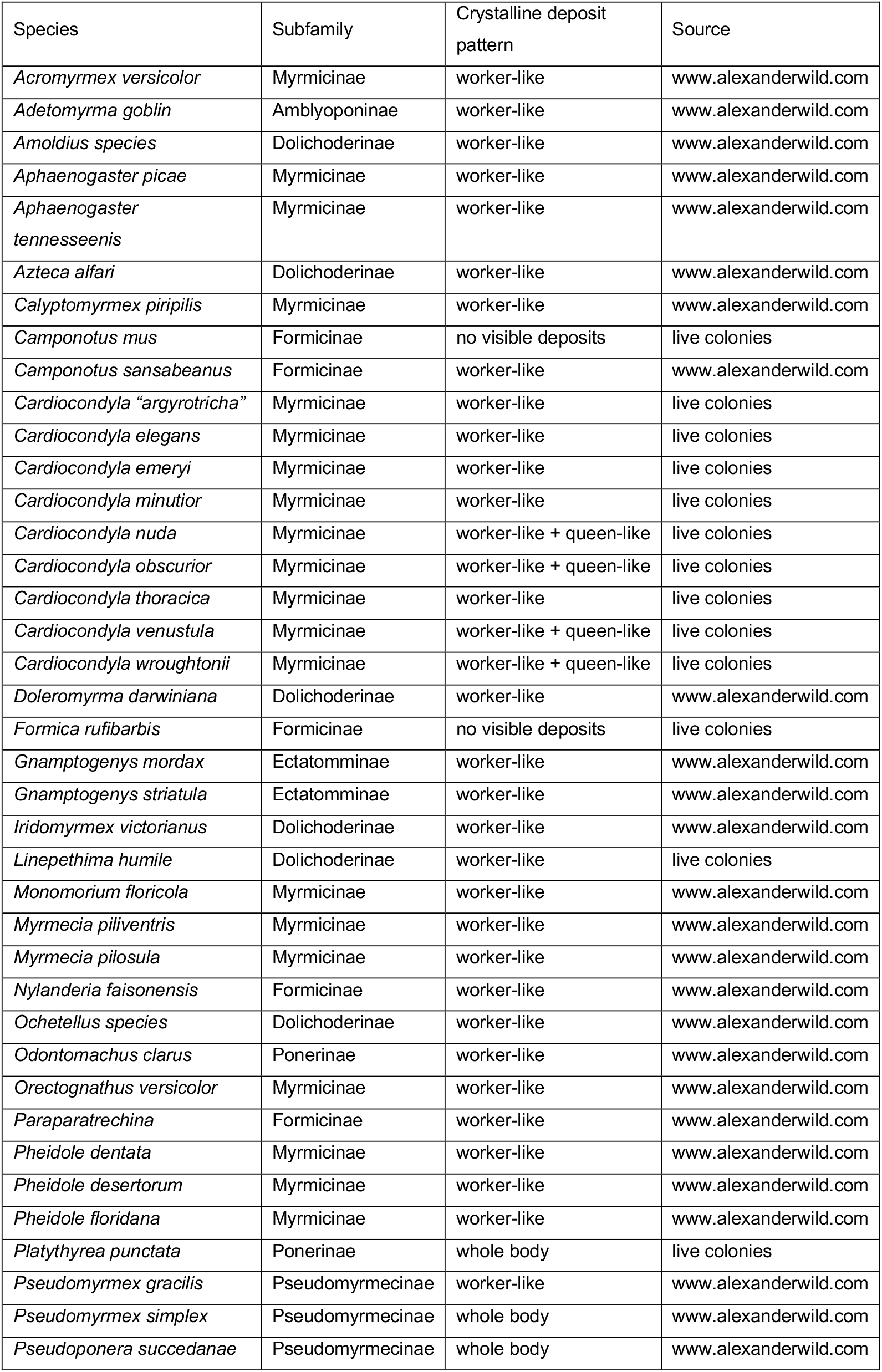
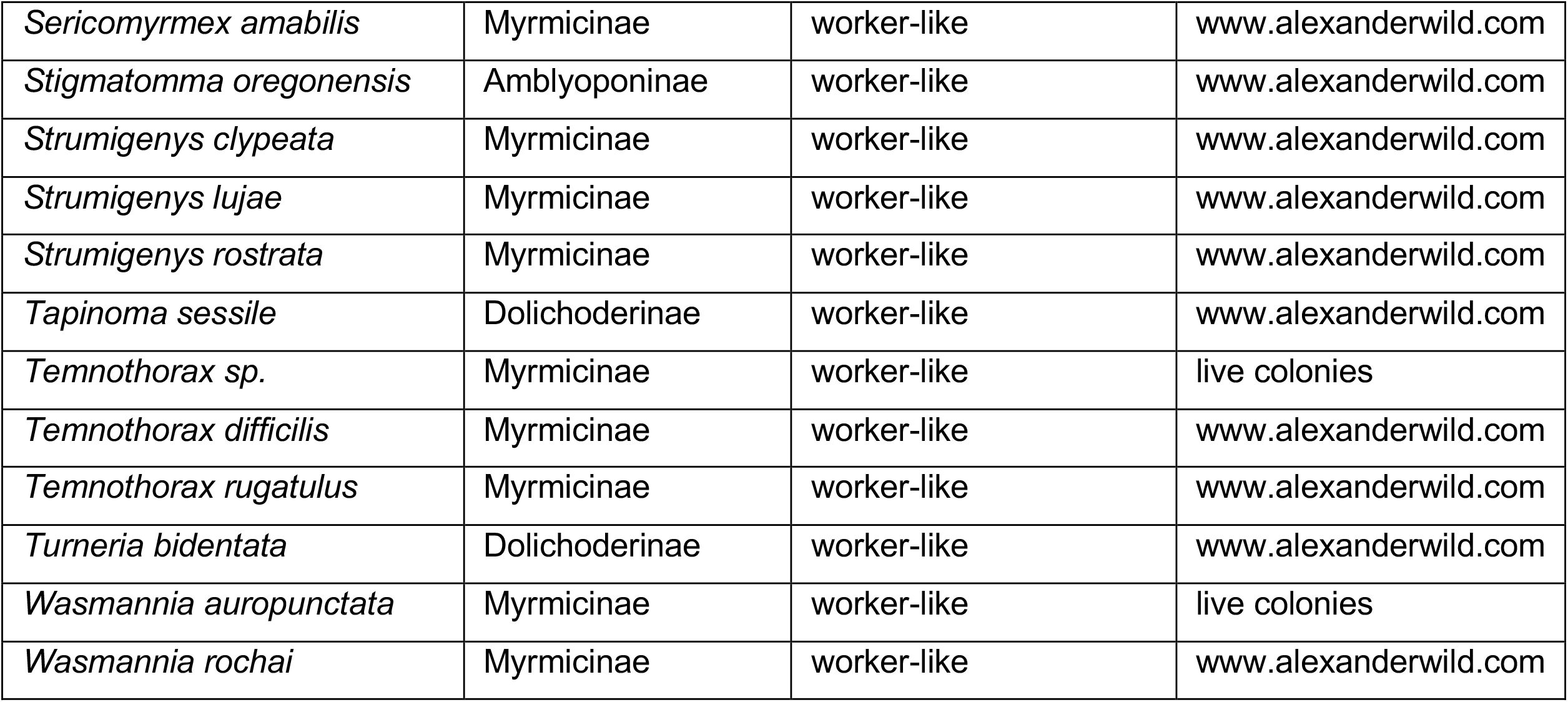
Overview of ant species screened for crystalline deposit patterns

